# A scalable approach for continuous time Markov models with covariates

**DOI:** 10.1101/2022.06.06.494953

**Authors:** Farhad Hatami, Alex Ocampo, Gordon Graham, Thomas E. Nichols, Habib Ganjgahi

## Abstract

Existing methods for fitting continuous time Markov models (CTMM) in the presence of covariates suffer from scalability issues due to high computational cost of matrix exponentials calculated for each observation. In this paper we propose an optimization technique for CTMM which uses a stochastic gradient descent algorithm combined with differentiation of the matrix exponential using a Padé approximation which we show then makes fitting large scale data feasible. We compare two methods for computing standard errors, one using Padé expansion, and the other using power series expansion of the matrix exponential. Through simulations we find improved performance relative to existing CTMM methods, and we demonstrate the method on the large-scale multiple sclerosis NO.MS dataset.

## 1 Introduction

Multiple sclerosis (MS) is a chronic neuroinflammatory and neurodegenerative disease that affects the central nervous system. Progression of the disease over time causes increasing disability, which is measured on the expanded disability status scale (EDSS) (Kurtzke [1983]), measuring several aspects of disability (such as walking) and indicating functioning ability of an individual, ranging between 0 (normal neurological examination) to 10 (death). As part of clinical trials, the EDSS score is an ordinal outcome measured approximately every 4 — 6 months as a part of clinical trials in relapsing and progressive MS patients. To understand better how the accumulation of disability is related to demographic and disease related variables, modeling of how the progression of EDSS varies from subject -to- subject, for example remains unchanged, increases (disease worsening) or decreases (disease improvement), is of interest. In addition, similar to many other neuroinflammatory and neurodegenerative diseases, MS is heterogeneous, i.e. patients’ EDSS scores change at individually varying rates, with a trend to gradually accumulate disability over the years.

A critical question for MS researchers is identifying the prognostic factors that influence EDSS progression. These factors provide insights to how progression varies between patients and point to possible directions for treatment. Different factors have been reported to be associated with the progression of disease; for instance, Vukusic and Confavreux [2007] showed age is a factor of progression, and Runmarker and Andersen [1993] showed 5 years after onset of MS, a low number of affected neurological systems, a low neurological deficit score and a high degree of remission from the last bout were the most important prognostic factors.

The analysis of disease progression in MS clinical trials has typically used time-to-event analysis, which is the recommended method for the demonstration of a drug effect on the first clinically meaningful disability progression in regulatory guidelines (Alvarez et al. [2015]). Other works on exploring prognostic factors in MS (Kappos et al. [2015], Vukusic and Confavreux [2003]) have have also relied on a time-to-event modeling framework. However, outside demonstrating a treatment effect on disease progression in randomized controlled trial settings, time-to-event models are not adequate for characterizing disease progression for three reasons; firstly, time-to-event models consider unidirectional changes and hence ignore that patients can improve or recover from their current EDSS state (see Mandel et al. [2007] for a detailed discussion), secondly, they do not account for repeated events of worsening thereby not fully using the available longitudinal information; and thirdly, these models cannot handle heterogeneity in EDSS transitions, i.e. time-to-event analysis treats worsening from EDSS 2 to 4 is the same as worsening from EDSS 4 to 6. However, factors that influence the early transitions (e.g. from EDSS 2 to 4) may not be the same as the factors influencing transitions at a later stage in the disease (e.g. from EDSS 4 to 6). For this reason, survival models for EDSS are subject to biases introduced by left censoring, i.e. they do not account for improvements in EDSS. Markov models address all of these concerns, allowing any transition (deterioration and recovery) and flexible modeling of covariates, where the covariate effect is estimated for each possible transition separately. A Markov process is one where the outcome at a given time only depends on the outcome of the previous time, and is independent of its past history. There are two types of Markov models; discrete and continuous. A discrete-time Markov model is one in which the system evolves through discrete time steps, while in a continuous-time Markov model, transitions between states can happen at any time. MS patients often have measurements that are taken at variable timepoints because in addition to regular check-ups, EDSS assessments are also conducted when patients relapse. An MS relapse is an acute neurological deterioration caused by a demyelinating event in the central nervous system, or in other words, a sudden worsening in MS symptoms, from which patients may or may not fully recover. Since EDSS measurements are not generated spontaneously at the exact time that a transition happens, therefore continuous time Markov model (CTMM) is a suitable model for such EDSS transition data. CTMM can generate the probability of moving from one EDSS state to another at any time, allowing one to estimate how factors are associated with faster or slower EDSS transition. CTMM are widely utilized to fit multi-state longitudinal data. These models in particular are applied in the field of public health, where the states of a Markov chain may refer to worsening stages of a chronic disease, such as breast cancer (Hsieh et al. [2002]. In longitudinal settings individuals are followed at regular intervals, where the exact transition times between disease states are typically not observed. Because state transitions can happen at any time and data is collected in a non-equidistant longitudinal manner, CTMM offer a more parsimonious approach over the discrete version of Markov models, and handle underlying heterogeneity nature of the disease.

A recent study (Lublin et al. [2022]) showed that relapse and age are important factors in the accumulation of disability in MS disease, although there is continued interest in which clinical factors are associated with EDSS transitions. Therefore, our goal here is to estimate the effect of different covariates on different EDSS transitions by applying CTMM on a large longitudinal dataset of MS patients. We are motivated by the large-scale NO.MS dataset, which includes approximately 20, 000 MS patients with longitudinal EDSS data followed for up to 15 years (Dahlke et al. [2021]), and with a range of clinically relevant covariates. However, the large sample size and serial measurements present a challenge for a CTMM because the likelihood and score function involves the evaluation of a matrix exponential for each observation, and as a result, model fitting and parameter estimation become burdensome, requiring numerical approximation methods (Moler and Van Loan [2003], Kalbfleisch and Lawless [1985]). Previously published papers focus on models in which the number of EDSS states have between restricted, with three or fewer state transitions allowed (Mandel et al. [2013]). However, in this paper we relax this assumption and allow for more granular transitions between different EDSS states. With a large database, and can thus explore the influence of covariates over these transitions. Mandel et al. [2007] developed a method for analyzing MS disease states using fixed-effects transition models, which along with their first-order Markov assumption suffered from lack of fit, partially due to the heterogeneous nature of the disease. NO.MS contains a large number of observations, allowing for more granular transitions between EDSS states, which results in a large number of model parameters, and the analysis becomes complex and computationally intense. In this paper, we propose a mini-batch stochastic gradient descent optimization technique combined with differentiation of the matrix exponential, based on the approach presented in Van Loan [1978] and Wilcox [1967]. This method has been previously applied to the context of hidden Markov models (Marshall and Jones [1995]), but to our knowledge has not been utilized for CTMM. We show that the mini-batch stochastic gradient used in this paper can accommodate large scale data. While other studies have modeled the transitions between different EDSS states without investigating clinical prognostic factors (Zurawski et al. [2019]), these factors or covariates affect the transition intensities between the different disease states. Indeed, the introduction of covariates should allow increased accuracy when predicting transition rates (Cook et al. [2002]). We show that our model can handle different types of covariates including baseline as well as time-varying covariates.

Section 2 represents the details of the model and inference. Section 3 presents our approach to estimation and inference in CTMM. Section 4 first describes how we simulate data from the proposed CTMM, and then describes how we assess performance of our models. Finally, Section 5 presents an application and discussion of the proposed model on the NO.MS.2 dataset.

## 2 Model

### 2.1 Continuous Time Markov Models (CTMM)

Consider a longitudinal study in which individuals can move among 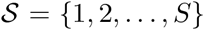 states. Let *M* denote the number of subjects in the study and *N_m_* denote the number of observation times for the mth subject. The time of subject *m*’s *k*th observation is denoted by *t_mk_*, when we observe the subject’s state as a random variable denoted by 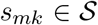. States are assumed to follow a first-order continuous time, discrete state Markov process, that is

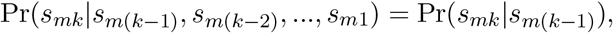

and we denote the probability of subject *m*’s transition from state *i* to state *j* as

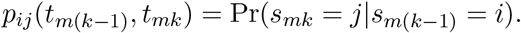

Figure 1 shows an illustration of state transitions and their corresponding probabilities. Individuals, after staying for some time in state *i*, move with some probability *p_ij_* from state *i* to state *j*. The likelihood for each individual is the product of all such transitions across observation times. The Markov process is fully characterized by its transition intensities

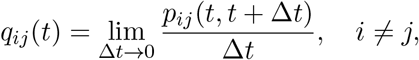

where 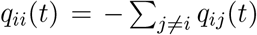 Cox and Miller [2017]. For the matrix representation, suppose **P**(*t*_1_,*t*_2_) and **Q**(*t*) denote the *S* × *S* matrix of transition probabilities *p_ij_*(*t*_1_,*t*_2_) and matrix of transition intensities *q_ij_* (*t*) respectively (the transition intensity matrix is also known as the rate or infinitesimal generator matrix). Each entry *q_ij_*(*t*) of the transition matrix **Q** represents the rate of transitioning from state *i* to state *j* at time *t*.

**Figure 1:**
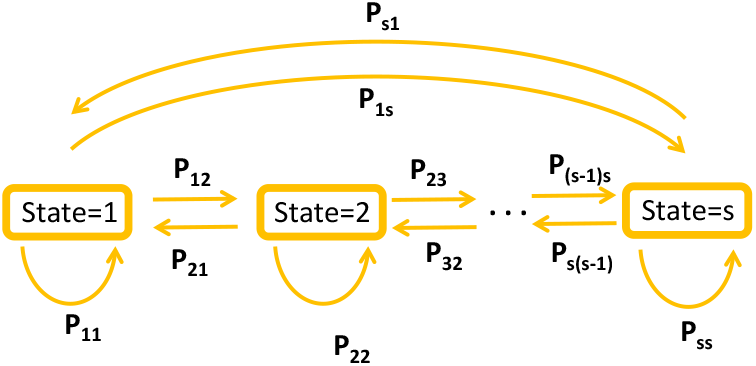
Transitions between states and their corresponding probabilities

Under time homogeneity, transition intensities are independent of the time i.e. *q_ij_*(*t*) = *q_ij_* and transition probabilities depend only on the elapsed time between successive observations i.e. *p_ij_*(*t*_1_;*t*_2_) = *p_ij_*(*t*_2_ – *t*_1_,0) for 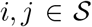. Then in this case, transition probabilities can be related directly to transition intensities through the so-called Kolmogorov equation,

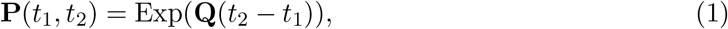

where Exp(·) indicates the matrix exponential (we denote the scalar exponential with exp(·)).

#### 2.1.1 Incorporating covariates in the model

Suppose subject *m* has *R* covariates *z_mr_*(*t_mk_*), *r* = 1,..., *R*, measured at time *t_mk_* (*k*-th observation), and written as the *R*-vector *Z_m_*(*t_mk_*). This could contain different types of covariates such as time-independent covariates (e.g. sex), as well as time-varying ones (e.g. age). We further assume that the covariate value remains constant into the future from the present value until the next observation. Marshall and Jones [1995] described a form of proportional hazards model, where in the presence of covariates, the transition intensity matrix elements *q_ij_* are replaced by

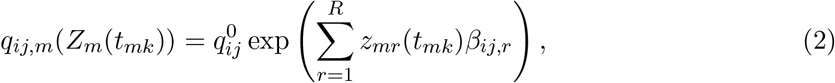

where *β_ij,r_* is a regression coefficients that quantifies the impact of each covariate on state transition from *i* to *j*, and 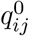 is the baseline intensity; we write *β* for the R × S × (S — 1)-vector of all coefficients, and **Q**^0^ is called baseline intensity matrix with entries 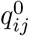. The interpretability of **Q**^0^ is the transition intensity when all covariates take on the value zero or at the reference value for categorical covariates, and as a result all covariates should generally be centered. We call this version of our model a “transition-dependent regression” where the effect of each covariate can vary for each state transition. We also can consider another version where regressors have a common effect over all state transitions- i.e. 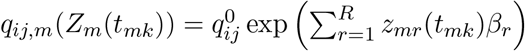, which we call the transition-independent regression case. For the rest of the paper, we will focus on the transition-dependent regression, though the transition-independent regression is modeled similarly and is nested within the more flexible model.

Let *τ_mk_* = *t_mk_* — *t*_*m*(*k*-1)_, *k* = 1, ⋯, *N_m_*, be the time lag between two consecutive observation times. Denote the parameters of interest *θ*, elements of **Q**^0^ and *β*, with dimension *R* × *S* × (*S* — 1). This allows us to formulate the time-invariant likelihood for subject *m* as follows

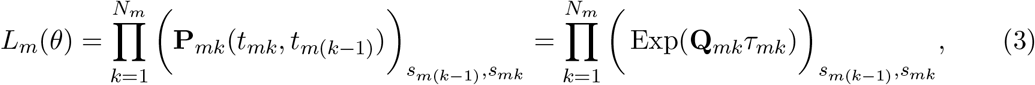

where (·)_*S*_*m*(*k*-1)_)*S*_*mk*__ refers to the corresponding entry of the matrix (row: state at time *t*_*m*(*k*-1)_, and column: state at time *t_mk_*). The complete likelihood then becomes product over all subjects, i.e. 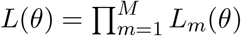. Our likelihood in (3) is the generalized version of likelihood used by [Kalbfleisch and Lawless, 1985], when covariate vectors are measured at every observation time (canonical decomposition of **Q** at every observation).

The ML estimator of *θ* can be obtained through the maximization of *L*(*θ*). Although *L*(*θ*) has a simple form, evaluation is computationally intensive because it includes the matrix exponential operation for each product of the likelihood. In the next section we introduce our approach to calculating the derivatives of *L*(*θ*) and develop a stochastic gradient descent approach to find the ML estimates.

## 3 Estimation & Inference

### 3.1 Optimization with Stochastic Gradient Descent

Combined together, Wilcox [1967] and Van Loan [1978] (Theorem 1) show that for any matrix function **A**(λ) = (*a_ij_*(λ)) for scalar λ, the derivative of Exp(**A**(λ)) w.r.t λ can be found via a Pade approximation

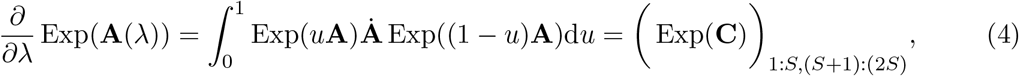

where 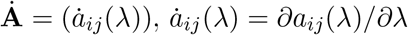 (i.e. element-wise derivative), 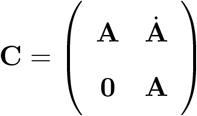, **0** is *S* × *S* zero matrix, and subscripts (·)_1:*S*,(*S*+1):(2*S*)_ indicate extraction of the upper right *S* × *S* submatrix.

Our goal is to obtain the partial derivatives of *L*(*θ*) w.r.t elements of *θ*, with *θ_ℓ_* standing in for each 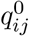 and *β_ij,r_*, for all transitions 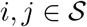 such that *i* ≠ *j* and all covariates *r*; for subject *m*

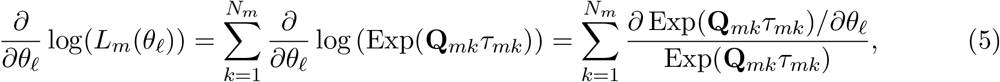

where the final ratio of matrices is evaluated as an entry-wise ratio for each entry (*i, j*), *i* ≠ *j*. Thus we need to calculate the derivatives for every subject *m* and at every observation time lag *τ_mk_*. To use the result (4) above, for each event *τ_mk_*, we take **A** = **Q**_*mk*_*τ_mk_* which gives 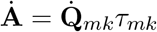, the partial derivatives w.r.t 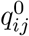,

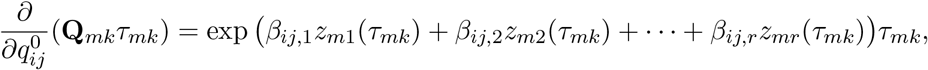

and elements corresponding to derivatives w.r.t *β_ij,r_*

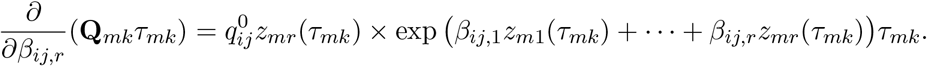

According to (4), having the matrices **Ȧ** for each observation time lag, we can form the block matrices **C** and use (5) to calculate *∂/∂θ* log(*L_m_*(*θ*)).

Finally, the gradient is obtained,

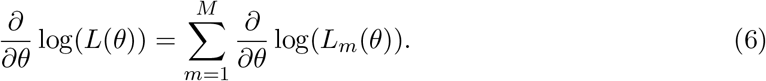

Having *∂/∂θ* log(*L*(*θ*)), we can now use gradient descent to find ML estimates of the parameters *θ* in (3). However, the standard gradient descent requires application on the full dataset, hence, computing matrix exponentials and derivatives at each observation time lag *τ_mk_* for all individuals. Consequently, it is computationally expensive to scale it to a large dataset. Instead, we use a mini-batch stochastic version of gradient descent Sakrison [1965], which randomly partitions the data into mini-batches and performs the calculations on each mini-batch instead of the entire dataset. This allows our method to scale to essentially arbitrarily large datasets. Let 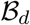 be a randomly chosen subset of subjects (with or without replacement) at the iteration step 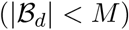. The updated parameters at step *d* + 1 of the mini-batch stochastic gradient descent are obtained via

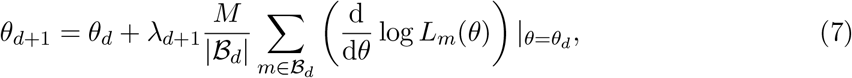

where λ_*d*+1_ is the learning rate sequence (we use decreasing learning rate with starting value equal to λ_*d*+1_ = (*d* + 1)^-0.6^). This procedure iterates until the parameters using (7) converge to 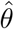 (i.e. 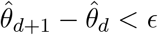 for a small *ϵ*). This then results in ML estimators of 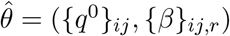, for all transitions 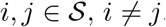, and all covariates *r*.

#### 3.1.1 Calculation of Confidence Intervals

Asymptotic standard errors and confidence intervals are computed from the Hessian matrix of the log likelihood evaluated at the parameter estimates. There are different numerical approximations/approaches in the literature to compute the Hessian for the parameters *θ*. Since calculation of the Hessian matrix requires computing (*S* × (*S* – 1) × *R*)^2^ elements, existing methods (Hanks [2018], Fleming and Calabrese [2021], and Jackson [2011a]) are computationally prohibitive if *M* is large. Here we propose two scalable approaches to calculate the Hessian matrix; the first is via calculation of second order derivatives through re-application of the Padé approximation in (4); and the second approach is with a second order approximation using the power series definition of the matrix exponential (see Moler and Van Loan [2003]).

##### 3.1.1.1 Padé expansion for Hessian

Let *θ* = (*θ*_1_,*θ*_2_) where *θ*_1_ and *θ*_2_ are the {*q*^0^}_*ij*_ and {*β*}_*j*_ components. The Hessian then has a 2 × 2 block form, with blocks (*ℓ,ℓ*’), with *ℓ,ℓ*’ ∈ {1, 2}. To obtain the Hessian matrix we need to calculate second derivative of the log-likelihood stated in (3) as follows

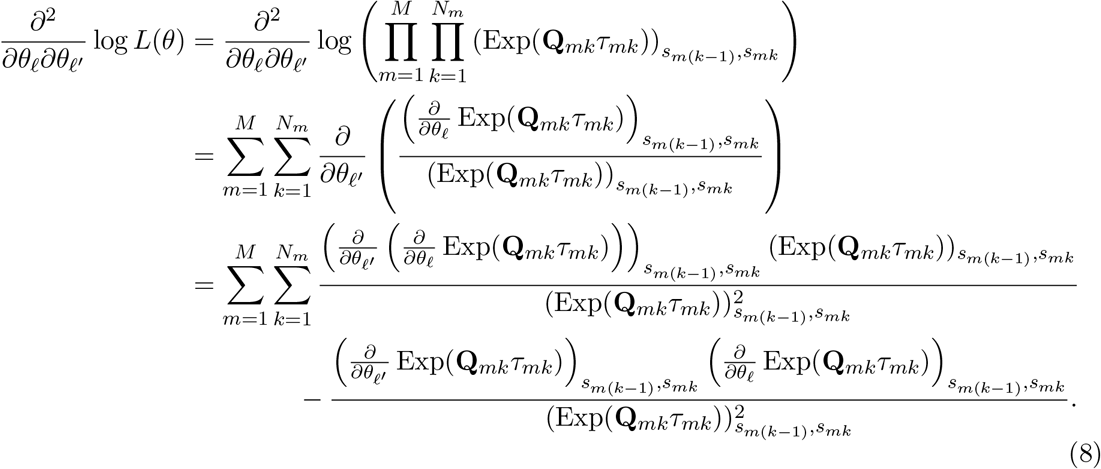

The terms in expression (8) can be found in Section 3.1, except for 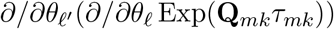, which is found as follows

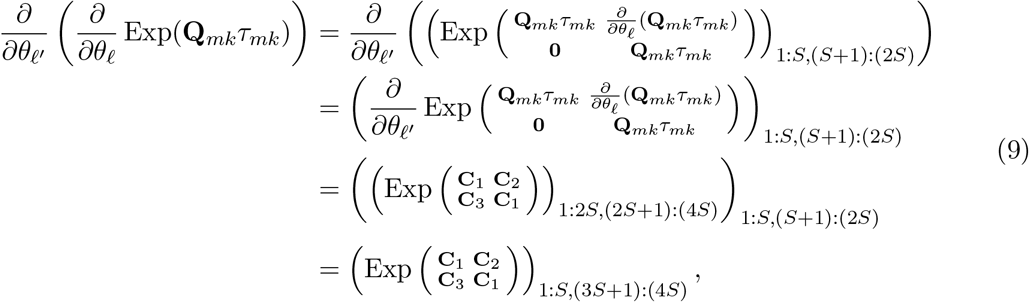

where 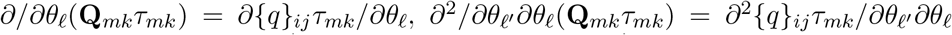, 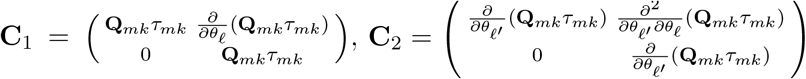,, **C**_3_ is 2*S* × 2*S* zero matrix, and **0** is an *S* × *S* zero matrix. Specification of the Hessian is completed by considering the different possible values for *θ*_*ℓ*’_ and *θ_ℓ_*:

1. 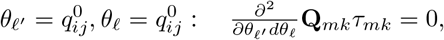
2. 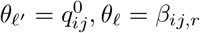 or 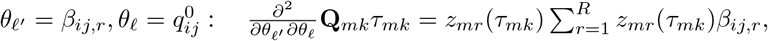
3. 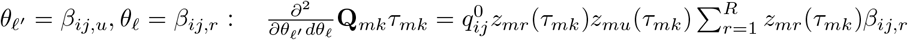

##### 3.1.1.2 Power series expansion for Hessian

Using the Padé approximation to calculate confidence intervals is computationally expensive especially when number of states *S* is large, due to the costly computation of the Exp function.

Thus we also introduce a second approach using approximation, which could be faster in terms of computation but with a slight compromise in accuracy. Recalling (3) and using the definition of matrix exponential we have

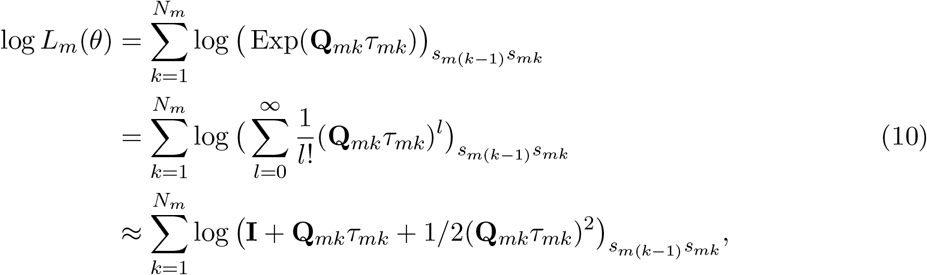

where **I** is the identity matrix, *s_mk_* is the occupied state at time *t_mk_*, and we truncate the power series after the third term (see Appendix for more details). We need to mention that the trade-off in this method is that truncation leads to slight underestimation of the variance in estimation of confidence intervals since some of the terms in the power series are dropped. We will explore the impact on coverage in section 4. Having the second derivatives we can simply form the Hessian matrix and calculation of confidence intervals then becomes straightforward.

### 3.2 Evaluation methods

We compare our method to an existing and widely used CTMM tool. Jackson [2011b] proposed a method that allows CTMM to be fitted to longitudinal data, and developed its R package called MSM as well (Jackson [2011a]). MSM uses different approaches for the optimization step, and we will evaluate each individually:

- MSM_opt: optim method which uses the deterministic Nealder-mead approach R Core Team [2021].
- MSM_nlm: Which uses Newton-type algorithm R Core Team [2021].
- MSMF: Which uses fisher scoring Kalbfleisch and Lawless [1985].

## 4 Simulation study

### 4.1 Simulation overview

We perform a simulation study under different scenarios to assess the performance of both the proposed methods for optimization by SCTMM and construction of confidence intervals outlined in section 3. Let *t_max_* be the maximum time in which subjects are followed up (for instance 10 years). Simulating data for the CTMM for one subject (m) is carried out via following steps:

1. We first choose an arbitrary baseline transition matrix **Q**^0^ and some values for the baseline covariates *z_mr_*(*t*_*m*0_), for every covariate *r*, then we can compute **Q**_*m*0_ via equation (2). The interpretation of the **Q**^0^ depends on the centering of the covariates. To obtain the stationary (also called steady) state probability (shown by π) we solve the following matrix equation:

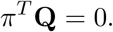 This specifies the initial state occupied by subject *m*. In other words, we draw a sample state from the finite space of states 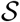 with probabilities *π*.
2. To obtain the time of the next state transition *t*_*m*(*k*+1)_, we draw one sample from an exponential distribution with rate — *q_S_mk_S_mk__*, where *s_mk_* stands for the state occupied by subject *m* at observation *k* (note that for the very first step of the simulation algorithm *k* = 0). Furthermore, randomly generate some values from uniform distribution for *z_mr_* (*t*_*m*(*k*+1)_) for every *r*.
3. Notice that the time *t*_*m*(*k*+1)_ generated at step 2, is the instantaneous transition time where subject *m* moves to any other state but not the currently occupied one *s_mk_*. Because of this, in order to simulate patients who remain clinically stable, we need to generate a number of dummy observations in which subject m has been observed several times staying at the same state but with different values of covariates (randomly). The choice of how many dummy sojourn observations would be required is arbitrary, however, it is advised to use the frequency in which the states are observed in the real dataset at hand to mimic the data; for instance, every 4 months.
4. Now we choose the next state *s*_*m*(*k*+1)_ by drawing a sample from the set of states 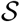 with probabilities computed by equation (1).
5. If *t*_*m*(*k*+1)_ < *t_max_* go to step 2, otherwise terminate the algorithm.

We then repeat the above steps for any given number of subjects *M*.

### 4.2 Simulation

We consider two main simulation scenarios; first a null case where there is no effect of covariates on the transition rates (i.e. *β_ij,r_* = 0 for every *i, j* and *r*); and a second case where there is an effect from the covariate impacting transition rates. In both cases we try to mimic the real dataset that we will use in the next section, and choose the number of states based on where we have the majority of transition data, and so we reduce the 20 EDSS states down to 8 states 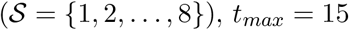 years (follow-up time). For the second case where the covariates impact the transition rates, we use 10 covariates and assess estimation of transition-dependent regression parameters *β_ij,r_*. All 10 covariates are randomly drawn from a uniform distribution and then centered at zero. We perform a sensitivity analysis over sample size *M* in both scenarios, and assess the performance of our SCTMM method. For both scenarios, and each sample size, we perform Monte Carlo simulation and generate 1000 realizations of the dataset, and evaluate the operating characteristics of bias, standard error, coverage, and rejection rates. Estimation with each realization uses different initial values for *β_ij,r_* for each *i, j*, and *r*; For each realization *β_ij,r_* are drawn from a normal distribution with mean 0 and standard deviation 1. The bench-marking platform used for this study ran R-4.0.0 to generate datasets and perform the analysis on 7 Intel Ivy Bridge cores each running at 1.15GHz speed and 16GB of RAM memory in total.

There are many configuration choices in implementing a mini-batch stochastic gradient optimization, and we take guidance from the existing literature (Franchini et al. [2020], Perrone et al. [2019]). With large scale data (*M* > 1000) it is advised to set the mini-batch size 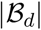 to be between 500 and 1000. In this paper we fix 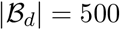.

The CTMM likelihood is not globally concave and thus multiple re-starts are recommended to get as close to the global maximum as possible; the optimization result that produces the largest log likelihood is taken as the optimal solution. Thus, we advise using between 1000 — 5000 re-starts with random initial values, for any number of parameters up to what we have here (we use 1000 re-starts). Careful selection of initial values is required to reduce the chance of the optimizer arriving at saddle points and local optimal. We suggest one of the following policies where applicable to set the starting parameters for the SCTMM method:

- Apply SCTMM on a smaller section of the data with no stochastic computation (for instance *M* = 500), estimate the parameters and then set these derived parameters for the restarts in the SCTMM.
- Sample off-diagonal elements for row *i* and column *j* from positive values of a univariate normal distribution with mean and standard deviation 1/|*i* – *j*|.

The above suggestions aim at minimizing the number of re-starts leading to highly suboptimal solutions and improve the accuracy of the estimates. We perform two analyses, one with no covariates in the model, where we estimate the transition intensities across all state transitions, and a second case where we include covariates and remove from the model any state transition *ij* that has less than 1% transition data in the entire dataset. Then we estimate covariate effects for this case. We also use the Padé expansion method to calculate confidence intervals.

### 4.3 Results

To assess the estimation performance of the coefficients *β_ij,r_* in the presence of covariate effects (i.e. the transition-dependent scenario) we have tested the performance of SCTMM against MSM. Table 1 shows a comparison between the optimization methods used in MSM to obtain maximum likelihood estimates, and our proposed SCTMM method. These results show that under the small-scale setting (*M* ≤ 1000) SCTMM has equal or better performance to MSM in terms of bias, variance, coverage and rejection rate. While all methods have some under coverage and slight inflation of the null rejection rate, while SCTMM is closest to the nominal. In a large-scale setting (*M* ≥ 10000) none of MSM’s methods were available (due to numerical instability) but the SCTMM approach estimates the parameters with reasonably good performance. Furthermore, the results demonstrate the scalability of the SCTMM approach to large datasets. Table 2 shows a comparison between the optimization methods used in the MSM software and our proposed SCTMM approach when there is no effect of any covariates incorporated in the model (*β_ij,r_* = 0 for every *i, j*, and *r*). Again, the results show that under a small-scale setting SCTMM has equal or better performance to MSM, and in a large-scale setting the SCTMM can handle the scalability problem. Figure 2 shows a comparison of the computation time between the SCTMM and MSM, for only one initialization (no re-starts), demonstrating that both methods are on-par in this case.

**Table 1:** Comparison of bias, variance, coverage and rejection rate between the methods used in the MSM software and our proposed SCTMM method, when varying *β_ij,r_*, sample size *M* and number of observations *N*.

| Transition-dependent |  |  |  |  |  |  |  |  |  |  |  |  |  |  |  |  |  |  |
| --- | --- | --- | --- | --- | --- | --- | --- | --- | --- | --- | --- | --- | --- | --- | --- | --- | --- | --- |
| M | N | True $\beta_{ij,r}$ | Bias | | | | Variance | | | | Coverage | | | | Rejection rate | | | |
|  |  |  | MSM_opt | MSM_nlm | MSM_F | SCTMM | MSM_opt | MSM_nlm | MSM_F | SCTMM | MSM_opt | MSM_nlm | MSM_F | SCTMM | MSM_opt | MSM_nlm | MSM_F | SCTMM |
| 500 | 15,663 | 0 | 0.01 | 0.12 | 0.011 | 0.009 | 0.006 | 0.008 | 0.006 | 0.004 | 0.928 | 0.9 | 0.920 | 0.932 | 0.072 | 0.075 | 0.073 | 0.068 |
| 500 | 15,663 | 0.5 | 0.012 | 0.14 | 0.12 | 0.01 | 0.009 | 0.014 | 0.01 | 0.009 | 0.93 | 0.9 | 0.93 | 0.94 | 0.71 | 0.72 | 0.70 | 0.70 |
| 500 | 15,663 | 2 | 0.022 | 0.23 | 0.022 | 0.015 | 0.009 | 0.02 | 0.01 | 0.01 | 0.92 | 0.91 | 0.89 | 0.93 | 0.81 | 0.8 | 0.81 | 0.83 |
| 1000 | 29,891 | 0 | 0.01 | 0.12 | 0.11 | 0.009 | 1e-5 | 1e-5 | 1e-4 | 1e-5 | 0.929 | 0.91 | 0.921 | 0.936 | 0.071 | 0.08 | 0.073 | 0.064 |
| 1000 | 29,891 | 0.5 | 0.005 | 0.01 | 0.009 | 0.004 | 0.009 | 0.013 | 0.009 | 0.009 | 0.93 | 0.89 | 0.9 | 0.94 | 0.71 | 0.68 | 0.69 | 0.70 |
| 1000 | 29,891 | 2 | 0.01 | 0.015 | 0.01 | 0.008 | 0.009 | 0.016 | 0.009 | 0.01 | 0.92 | 0.88 | 0.91 | 0.92 | 0.80 | 0.79 | 0.80 | 0.82 |
| 10,000 | 312,547 | 0 |  |  |  | 0.016 |  |  |  | 0.005 |  |  |  | 0.91 |  |  |  | 0.09 |
| 10,000 | 312,547 | 0.5 |  |  |  | 0.018 |  |  |  | 0.01 |  |  |  | 0.94 |  |  |  | 0.72 |
| 10,000 | 312,547 | 2 |  |  |  | 0.016 |  |  |  | 0.009 |  |  |  | 0.93 |  |  |  | 0.83 |
| 20,000 | 615,720 | 0 |  |  |  | 0.015 |  |  |  | 0.01 |  |  |  | 0.92 |  |  |  | 0.08 |
| 20,000 | 615,720 | 0.5 |  |  |  | 0.019 |  |  |  | 0.02 |  |  |  | 0.91 |  |  |  | 0.73 |
| 20,000 | 615,720 | 2 |  |  |  | 0.017 |  |  |  | 0.013 |  |  |  | 0.90 |  |  |  | 0.81 |

**Table 2:** Comparison of bias, variance, coverage and rejection rate, in the setting where there are no covariates, between the methods used in the MSM software and our proposed SCTMM method, when varying 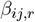, sample size *M* and number of observations *N*.

| Estimation of baseline intensities in absence of covariates ($\beta_{ij,r} = 0$) | | | | | | | | | | | | | | | | | | |
| --- | --- | --- | --- | --- | --- | --- | --- | --- | --- | --- | --- | --- | --- | --- | --- | --- | --- | --- |
| M | N | True $\tilde{q}_{ij}^0$ | Bias | | | | Variance | | | | Coverage | | | | Rejection rate | | | |
|  |  |  | MSM_opt | MSM_nlm | MSM_F | SCTMM | MSM_opt | MSM_nlm | MSM_F | SCTMM | MSM_opt | MSM_nlm | MSM_F | SCTMM | MSM_opt | MSM_nlm | MSM_F | SCTMM |
| 500 | 15,663 | 0 | 7e-6 | 8e-6 | 7e-6 | 7e-6 | 5e-6 | 7e-6 | 5e-6 | 6e-6 | 0.952 | 0.949 | 0.952 | 0.953 | 0.01 | 0.05 | 0.01 | 0.01 |
| 500 | 15,663 | 0.5 | 9e-6 | 2e-5 | 9e-6 | 9e-6 | 7e-6 | 9e-6 | 7e-6 | 7e-6 | 0.951 | 0.949 | 0.952 | 0.951 | 0.73 | 0.71 | 0.72 | 0.73 |
| 500 | 15,663 | 1 | 1e-5 | 3e-5 | 4e-5 | 1e-5 | 7e-6 | 1e-6 | 7e-6 | 7e-6 | 0.94 | 0.939 | 0.941 | 0.94 | 0.84 | 0.8 | 0.83 | 0.89 |
| 1000 | 29,891 | 0 | 2e-5 | 4e-5 | 2e-5 | 1e-5 | 9e-6 | 2e-5 | 9e-6 | 8e-6 | 0.947 | 0.94 | 0.948 | 0.949 | 0.08 | 0.1 | 0.09 | 0.01 |
| 1000 | 29,891 | 0.5 | 4e-5 | 6e-5 | 4e-5 | 3e-5 | 3e-5 | 8e-5 | 2e-5 | 2e-5 | 0.949 | 0.94 | 0.952 | 0.946 | 0.77 | 0.75 | 0.77 | 0.79 |
| 1000 | 29,891 | 1 | 5e-5 | 7e-5 | 4e-5 | 4e-5 | 8e-5 | 1e-4 | 8e-5 | 7e-5 | 0.936 | 0.931 | 0.935 | 0.941 | 0.81 | 0.73 | 0.82 | 0.87 |
| 10,000 | 312,547 | 0 |  |  |  | 0.001 |  |  |  | 5e-4 |  |  |  | 0.94 |  |  |  | 0.012 |
| 10,000 | 312,547 | 0.5 |  |  |  | 0.003 |  |  |  | 0.003 |  |  |  | 0.937 |  |  |  | 0.78 |
| 10,000 | 312,547 | 1 |  |  |  | 0.008 |  |  |  | 0.006 |  |  |  | 0.931 |  |  |  | 0.89 |
| 20,000 | 615,720 | 0 |  |  |  | 0.009 |  |  |  | 0.007 |  |  |  | 0.941 |  |  |  | 0.02 |
| 20,000 | 615,720 | 0.5 |  |  |  | 0.01 |  |  |  | 0.009 |  |  |  | 0.935 |  |  |  | 0.90 |
| 20,000 | 615,720 | 1 |  |  |  | 0.02 |  |  |  | 0.01 |  |  |  | 0.93 |  |  |  | 0.88 |

**Figure 2:**
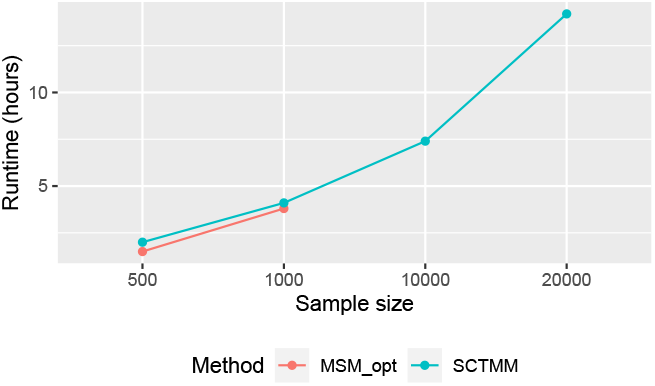
Running time comparison between MSM software and the proposed SCTMM method.

## 5 Application on NO.MS.2 dataset

### 5.1 NO.MS.2 data

We utilize the novel Novartis-Oxford MS (NO.MS.2) dataset in this paper, formed by aggregating multiple sclerosis patient data from 34 MS clinical trials and their extensions (Dahlke et al. [2021], Lublin et al. [2022]). Currently, NO.MS.2 is the largest and most comprehensive clinical trial dataset in MS, spanning all MS phenotypes and containing data from over 27, 000 patients with up to 15 years of follow-up visits, and with regular monitoring of patients’ neurological status by highly trained raters across all stages of MS disease. The earliest clinical trial in this dataset began in 2003 and the latest measurement is from January of 2020. These trials collected longitudinal data on disability, as measured by Kurtzke’s Expanded Disability Status Scale (EDSS) (Kurtzke [1983]). In this analysis EDSS was simplified as follows: state 0 remained 0; for EDSS 1 – 1.5, 2 – 2.5, 3 – 3.5, ⋯, were rounded down to 1, 2, 3, ⋯; EDSS? 8 was set to 8, resulting in 9 reduced states.

In this work we round half EDSS values to the next largest integer.

In this study we used a cohort of *M* = 13, 320 patients, with *N* = 170, 628 observations in total, *S* = {0,1 ⋯, 8} EDSS states and a set of *R* = 7 covariates including the followings:

- ARR1: The annualized relapse rate is the number of relapses a patient experiences within the year prior to the current observation time (time-varying).
- Age: Age at event time divided by 10, to interpret coefficients as the effect by decade of age and to make coefficients more comparable with other variables (time-varying).
- DURFS: Duration of MS in years from first symptoms as estimated at trial entry (baseline).
- MSTYPE: A 3 level categorical variable, indication the phenotype of MS disease, with levels relapsing-remitting MS RRMS (used as reference level), secondary progressive MS (SPMS), and primary progressive MS (PPMS).
- Sex: Male or female (reference level female).

We perform two analyses:

1. First, an analysis where we include no covariates in the model, and we consider all EDSS transitions (i.e. keeping rare transitions).
2. Second, an analysis where we only allow transitions that have more than 1% of transition data in the entire dataset, which for this particular dataset turns out to be mostly EDSS transitions with jumps of more than 2 states. For these transitions we fit covariate effects.

All covariates are centered and we apply SCTMM using mini-batches of size 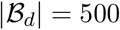 in each iteration step *d*. We also use 1000 restarts with different initial values of *θ*, and draw these initial values from a uniform distribution with min = 0, max = 1 and min = –1, max = 1 for {*q*^0^}_*ij*_ and {*β*}_*ij,r*_, respectively.

### 5.2 Results

Figure 3 (right side) shows the estimation of the baseline transition probability matrix when there is no covariate effects both in the data and model. It can be seen that patients are most likely to stay unchanged in their current states and then the probability of transitioning to a higher/lower EDSS state is higher in the early-mid stages of the disease than in later stages of the disease. The plausibility of these results has also been checked by looking at 3 (left side), a normalized empirical transition probability matrix which is conduced based on the transition data in the entire dataset.

**Figure 3:**
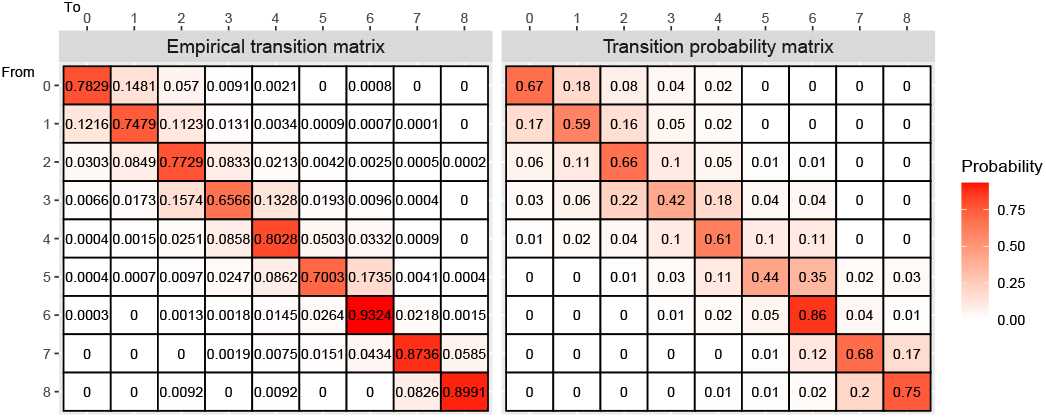
(Left) normalized empirical transition matrix. (Right) estimated transition probability matrix by SCTMM.

The graphs shown in Figure 4 illustrate transition-specific hazard ratios (exp(*z_r_β_ij,r_*)), when the covariate is set to its mean value, and corresponding 95% confidence intervals (all covariates are included in the model at the same time). These graphs show the effect of each covariate on successive EDSS transitions. Figure 4a demonstrates hazard ratios of ARR1, showing that relapses have a significant association on the accumulation of EDSS disability, particularly in but not limited to, the early stages of the disease. This is in concordance with the findings of a recent study by Lublin et al. [2022]. Based on these results, for instance, having a relapse in the past year for someone being at EDSS 3, will increase the risk of transition to EDSS 4 by approximately 40%(25% — 50%), controlling for other covariates. Figure 4b shows that older age is associated with faster EDSS transition, primarily but not exclusively early in the disease. For instance, having a decade increase in age increases the risk of transitioning from EDSS 2 to 3 by approximately 19%(14% — 24%). Figures 4c shows some association of duration of the MS disease since first symptoms were observed on the EDSS transitions.

**Figure 4:**
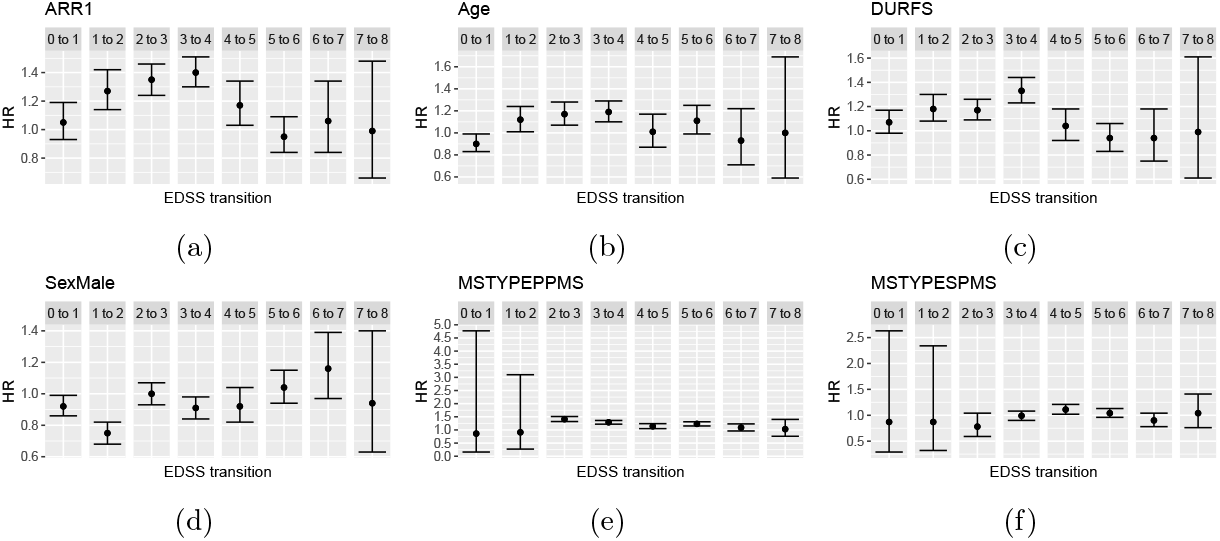
Estimated hazard ratios of different covariates on successive EDSS state worsening. Error bars show 95% confidence intervals.

Figures 5 show a more comprehensive heat-map plot of hazard ratios of different covariates, where we do not only show the successive transitions but more EDSS changes up to two jumps of states..

**Figure 5:**
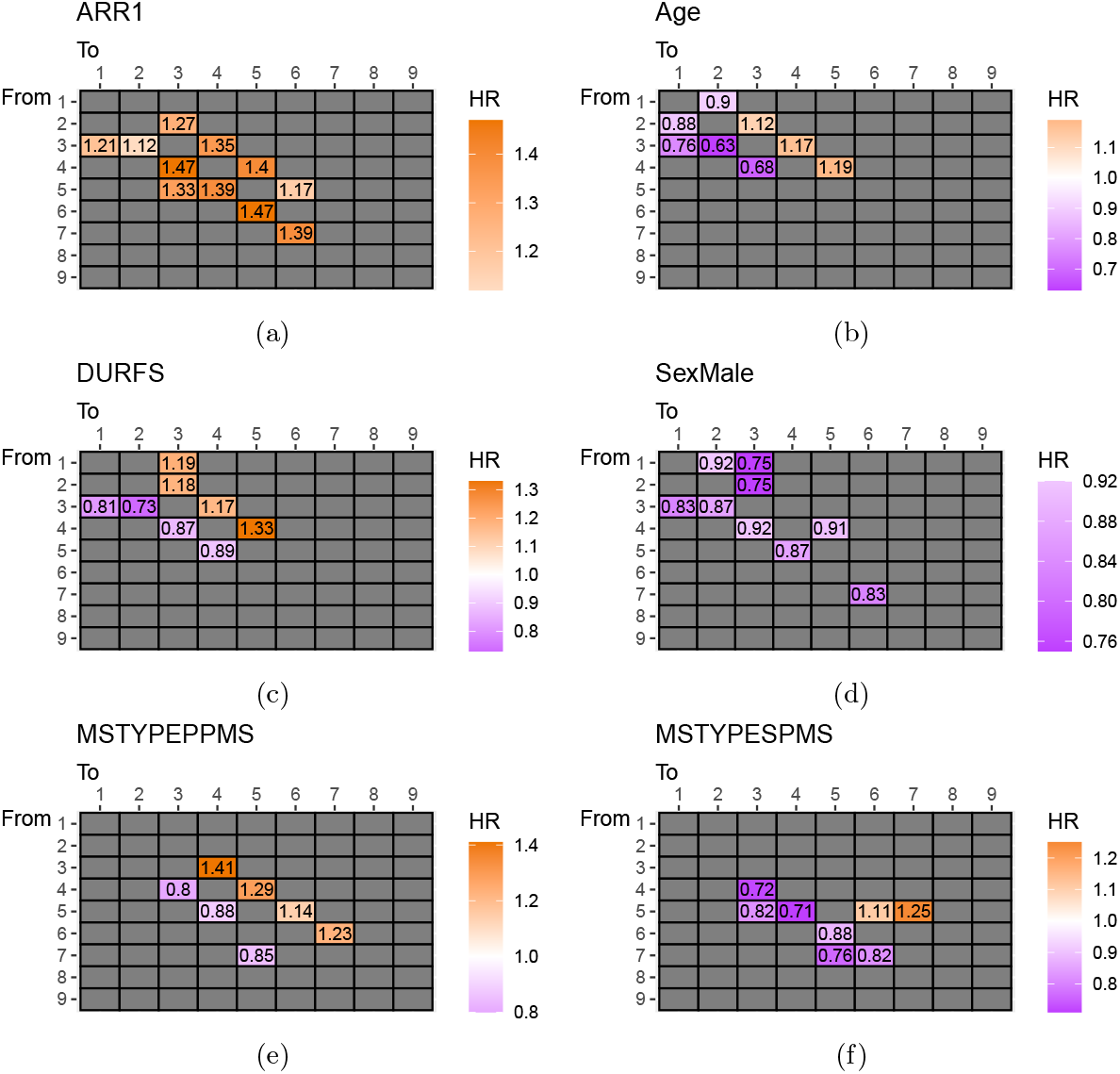
Estimated significant hazard ratios (HR 95% CI does not include 1) for different covariates on EDSS state transitions.

Figures 5a and 5b show that having a relapse in the last year and older age impact chance of EDSS worsening or improvement particularly at the early stage of the disease (most patients at this stage of the disease are of the ‘relapsing-remitting MS’ sub-type, hence the name). Figure 5c shows the longer the duration of disease, the higher the chance of further deterioration (HR> 1) and lower the chance of recovery (HR < 1), controlling for other covariates including age. It may be expected that covariates have opposing effect on the risk of worsening versus improvement, however, figures 5a and 5d show that this is not the case of having a relapse and being male; i.e. having a relapse in the last year always increase both chances of deterioration and improvement (e.g. recovery from relapses), and males have a lower hazard of transitioning compared to females (this may likely be explained by the higher probability of male patients belonging to the progressive subtypes where the female to male ratio is approximately 1:1 than the relapsing-remitting subtype of MS where the corresponding ratio may be 2:1 or even 3:1). Figures 5e and 5f show that patients with a diagnosis of progressive type of the disease (SPMS or PPMS) have a higher change of deterioration and a reduced change of recovery compared to patients diagnosed with RRMS.

## 6 Conclusion

We have proposed a method to overcome the problems associated with fitting CTMM with covariates to large-scale datasets. We use a mini-batch stochastic gradient descent algorithm which uses a smaller random subset of the dataset at each iteration, making it practical to fit large scale data. Furthermore, we used the results introduced by Wilcox [1967] and Van Loan [1978] to calculate the derivatives of the matrix exponential, and using this, then proposed a novel approach for computing confidence intervals via two applications of a Padé approximation to find the second derivatives. We also proposed another method for computing confidence intervals based on the approximation of the power series definition of the matrix exponential. The latter is useful when the number of states in the CTMM problem is high (≈ *S* > 20). In a small/mid scale setting (*M* ≤ 1000) our simulation studies show slight out-performance of the proposed SCTMM over the MSM software. In a large-scale setting (*M* ≥ 10,000), where the MSM software is unable to estimate the parameters, the proposed SCTMM can be used and shows a good performance. Some of the important findings in the analysis of NO.MS.2 are as follows:

- Applying the proposed SCTMM on the large scale NO.MS.2 dataset it is concluded that the number of relapses in the last year (ARR1) and older age increase the risk of deterioration (EDSS increase) and reduce the chance of recovery (EDSS decrease) particularly in the early stage of the disease.
- Disability accumulation is a slow process in MS: In our large MS dataset in which EDSS assessments occur on average approximately every 4 months, most patients stay unchanged in their EDSS state between consecutive visits. The probability of transitioning to a higher/lower EDSS state is occurring more frequently in the young and relapsing-remitting patient subtype compared with the progressive subtypes of the disease (SPMS or PPMS) where patients are more likely to worsen and less likely to recover.
- Our proposed statistical method makes the fitting of Markov models with covariates feasible and scalable to large datasets. It allows the investigation of covariate effects on transition probabilities between (disease) stages, which may find its application far beyond MS.

## Acknowledgement

The authors acknowledge the work of Jelena Cuklina, Steve Gardiner and Piet Aarden in coordinating and conducting data wrangling work. They also thank Dieter A. Häring for his detailed comments on the paper.

## Appendix

To compute the first derivatives w.r.t. 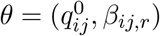, we need to only consider the *s*_*m*(*k*-1)_*s_mk_* entry of the matrix exponential:

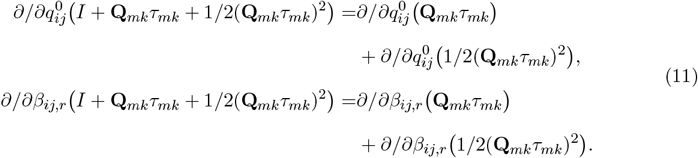

Now *∂/∂*(·)(**Q**_*mk*_*τ_mk_*} are as calculated in section 3, and we are left with calculation of *∂/∂*(·)(1/2(**Q**_*mk*_*τ_mk_*)^2^). To extract the entry *s*_*m*(*k*-1)_*s_mk_* we can use the general form for the second power of the matrix 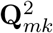,

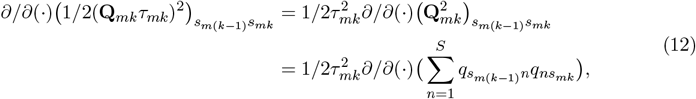

and the same goes for derivatives w.r.t *β_ij,r_*. Now define

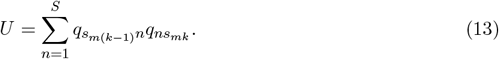

Applying the chain rule i.e. 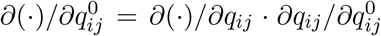 (and similarly for *β_ij,r_*) we can consider the three following situations:

1. If *i* – *s*_*m*(*k*-1)_ and *j* = *s_mk_*:

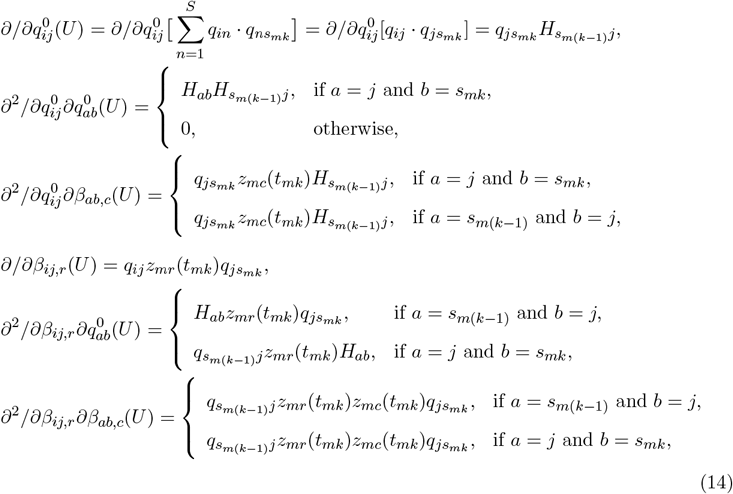

where 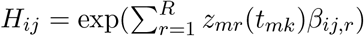.
2. If *i* = *s*_*m*(*k*-1)_ and *j* — *s_mk_*:

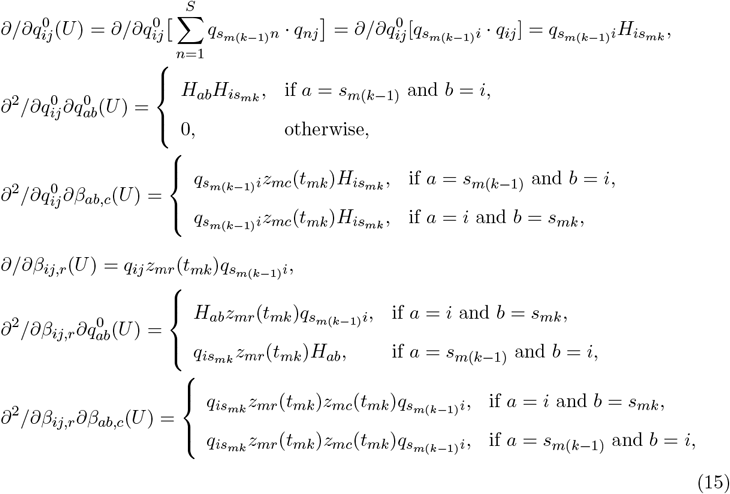
3. If *i* — *s*_*m*(*k*-1)_ and *j* – *S_mk_*:

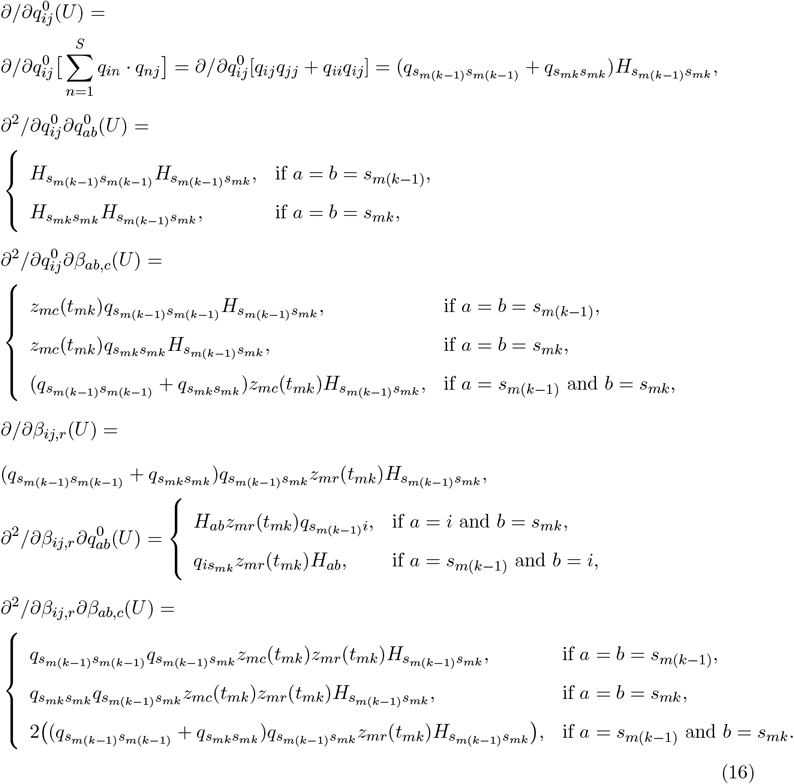 Notice that when *i* = *s*_*m*(*k*-1)_ and *j* = *s_mk_* then derivatives w.r.t both 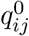 and *β_ij,r_* are zero.

